# A TRAIL-R4 splice isoform sensitizes cancer cells to TRAIL-induced apoptosis

**DOI:** 10.64898/2026.11.22.768413

**Authors:** Colin Poirier, Louis-Philippe Picard, Pierre Lavigne, Benoit Laurent

## Abstract

The immune system can recognize and eliminate tumor cells through the secretion of specific cytokines, including TRAIL (tumor necrosis factor-related apoptosis-inducing ligand). TRAIL triggers apoptosis by binding to its death receptors, TRAIL-R1 and TRAIL-R2. However, cancer cells can evade TRAIL-mediated cell death through the expression of decoy receptors, TRAIL-R3 and TRAIL-R4, which can bind TRAIL but lack the ability to initiate apoptotic signaling. Here, we focused on the decoy receptor TRAIL-R4, also known as TNFRSF10D, and investigated an alternative splicing event that generates a transcript lacking exon 3. This exon-skipping event results in a truncated receptor isoform, termed iTRAIL-R4, which lacks part of the ligand-binding domain. Structural biology analyses confirmed that TRAIL cannot efficiently bind to iTRAIL-R4. Interestingly, whereas the canonical TRAIL-R4 isoform protects cancer cells against TRAIL-induced apoptosis, overexpression of iTRAIL-R4 sensitizes cancer cells to TRAIL-mediated cell death. Furthermore, combining iTRAIL-R4 expression with different therapeutic treatments enhances the sensitivity of cancer cells. Collectively, our findings highlight the potential of iTRAIL-R4 to help counteract therapeutic cancer resistance and recurrence. They further support the therapeutic relevance of modulating TRAIL-R4 alternative splicing to shift cancer cells toward a more apoptosis-sensitive state.

## Introduction

TRAIL (TNF-related apoptosis-inducing ligand) is a cytokine involved in immune surveillance and the elimination of tumor cells (Falschlehner et al., 2009; Pimentel et al., 2023). Since its discovery (Ashkenazi et al., 1999a; Wiley et al., 1995), TRAIL has attracted considerable interest because of its ability to selectively induce apoptosis in tumor cells while largely sparing normal tissues (Ashkenazi et al., 1999b). This apparent therapeutic selectivity has prompted extensive investigation into TRAIL-based anticancer strategies. However, despite encouraging preclinical findings, clinical trials have yielded disappointing results, with limited or no significant therapeutic benefit in many patients (Montinaro & Walczak, 2022; Stuckey & Shah, 2013). These outcomes highlight the need for a better understanding of the molecular mechanisms governing TRAIL receptor activation and the cellular factors that determine sensitivity or resistance to TRAIL-induced apoptosis (Szegezdi, Van Der Sloot, et al., 2012; Thorburn et al., 2008).

TRAIL induces apoptosis through two death receptors, TRAIL-R1 and TRAIL-R210–13. Ligand binding promotes receptor oligomerization and the recruitment of the adaptor protein FADD, together with initiator caspases-8 and-10, leading to the formation of the death-inducing signaling complex (DISC) (Chinnaiyan et al., 1995; Mahmood & Shukla, 2010; Walczak & Sprick, 2001). Although both receptors activate similar downstream apoptotic pathways, they exhibit distinct activation properties depending on ligand presentation and receptor cross-linking (Kuijlen et al., 2010; McCarthy et al., 2006; von Karstedt et al., 2015). TRAIL-R1 is preferentially activated by membrane-bound or highly cross-linked TRAIL, whereas TRAIL-R2 can efficiently respond to soluble TRAIL. Following DISC formation, activated initiator caspases trigger the downstream apoptotic cascade.

In addition to these two death receptors, TRAIL interacts with two decoy receptors, TRAIL-R3 and TRAIL-R4. TRAIL-R3 lacks an intracellular domain and is therefore unable to transmit apoptotic signals (Labovsky et al., 2017; Mérino et al., 2006; Sheikh et al., 1999), whereas TRAIL-R4 contains a truncated death domain that is insufficient to initiate canonical apoptotic signaling (Degli-Esposti et al., 1997a; Marsters et al., 1997; Pan et al., 1998). Both receptors can bind TRAIL and have consequently been proposed to limit apoptosis by competing with TRAIL-R1 and TRAIL-R2 for ligand availability. However, TRAIL-R4 appears to exert more complex regulatory functions. Unlike other decoy receptors, it does not exhibit a higher affinity for TRAIL than the apoptotic receptors (Lee et al., 2005a; Truneh et al., 2000), raising questions about the mechanisms underlying its anti-apoptotic activity (Lang et al., 2016; Lee et al., 2005a; Wajant, 2019a). TRAIL-R4 may therefore inhibit apoptosis not only through ligand competition but also through interactions with other TRAIL receptors. Indeed, TRAIL-R1, TRAIL-R2, and TRAIL-R4 can form homo-and heteromeric receptor complexes, including ligand-independent associations (Lee et al., 2005a; Neumann et al., 2014; Wajant, 2019a). In particular, heterodimerization of TRAIL-R4 with the death receptors has been proposed to interfere with receptor organization and DISC assembly, thereby limiting initiator caspase activation.

The biological functions of TRAIL-R4 may extend beyond its ability to regulate apoptotic receptor signaling. Unlike TRAIL-R3, TRAIL-R4 retains a cytoplasmic region capable of activating non-apoptotic signaling pathways, including NF-κB (Ashkenazi et al., 1999a; Degli-Esposti et al., 1997a; Hu et al., 1999; Rambow et al., 2022; Yang et al., 2018; Zhou et al., 2017). However, the contribution of these pathways to tumor cell survival and progression remains poorly understood. Consistent with a potential role in tumor biology, TRAIL-R4 overexpression has been associated with poor prognosis and more aggressive tumor phenotypes in several cancers (Ganten et al., 2009; Sanlioglu et al., 2007). Conversely, reduced TRAIL-R4 expression has also been reported in other cancer types (Burns & El-Deiry, 2001; Daniels et al., 2005; Sanlioglu et al., 2005), suggesting that its biological significance may depend on cellular context. Moreover, TRAIL-R4 expression does not consistently predict cellular sensitivity to TRAIL. Elevated transcript levels have been reported in several TRAIL-sensitive tumor cell lines, including SW480, A549, and HeLa (Burns & El-Deiry, 2001; Degli-Esposti et al., 1997b; Xu et al., 2003). Together, these observations suggest that TRAIL-R4-mediated regulation of apoptosis involves mechanisms that cannot be explained solely by its expression level or ligand-binding properties.

Alternative splicing represents an additional layer of regulation that could contribute to the functional diversity of TRAIL receptors. Several splice variants have been described within this receptor family, including alternative isoforms of TRAIL-R2 (López-Gómez et al., 2016.; Screaton et al., 1997). In 2008, an additional TRAIL-R4 transcript, termed β-TRAIL-R4, was identified in renal cancer cells (Krieg et al., 2006). This isoform results from alternative splicing of exon 3 of the *TTRAIL-R4* gene (also known as *TNFRSF10D*) and has remained largely uncharacterized. Exon 3 skipping removes a portion of the extracellular region of TRAIL-R4, potentially affecting receptor organization, oligomerization, and ligand recognition. Consequently, this alternative splicing event could substantially alter the functional properties of TRAIL-R4. However, whether this isoform affects TRAIL binding, receptor-mediated signaling, or cellular sensitivity to TRAIL-induced apoptosis has not been established.

In this study, we investigated the structural and functional consequences of TRAIL-R4 alternative splicing, focusing on the exon 3-skipped isoform, which we termed iTRAIL-R4. We first examined how exon 3 skipping affects the extracellular architecture of the receptor and its predicted interactions with TRAIL. We then evaluated the respective effects of canonical TRAIL-R4 and iTRAIL-R4 on TRAIL-induced apoptotic signaling in MDA-MB-231 breast cancer cells, as well as the consequences of altering the relative expression of the two isoforms. Finally, we investigated whether iTRAIL-R4 expression could influence cellular responses to TRAIL-based combination treatments with conventional chemotherapeutic agents. By characterizing this previously unexplored TRAIL-R4 isoform, our study provides new insights into the regulation of TRAIL receptor signaling and identifies alternative splicing as a potential mechanism influencing tumor cell sensitivity to apoptosis.

## Methods

### Plasmid construction

The human TRAIL-R4/TNFRSF10D coding sequence was amplified from a human cancer cDNA library using Q5 High-Fidelity DNA Polymerase (New England Biolabs). Forward and reverse primers containing respectively NotI and AscI restriction sites, were used to amplify the full-length sequence. The stop codon was omitted to allow subsequent addition of a C-terminal epitope tag. Primer sequences are provided in **Supplementary table S1**. The cDNA encoding the TRAIL-R4 isoform (iTRAIL-R4) was generated using a synthetic 338-bp fragment comprising exons 1 and 2 and the first 70 nucleotides of exon 4, excluding exon 3. The fragment was amplified using the forward primer designed for the full-length construct and a reverse primer incorporating a HindIII restriction site. The remaining sequence, spanning the middle of exon 4 to exon 9, was amplified separately using a forward primer containing the HindIII site and the reverse primer used for the full-length construct. Amplified full-length TRAIL-R4 canonical and isoform were subsequently inserted to a pTOPO vector (ThermoFisher Scientific). The constructs were then transferred from the pTOPO cloning vector into a lentiviral destination vector (pHAGE lentiviral vector with a C-terminal FLAG and HA tag; pHAGE-C-FLAG-HA) using Gateway LR Clonase (ThermoFisher Scientific), according to the manufacturer’s instructions.

### Cell culture

Human embryonic kidney HEK-293T cells and human breast adenocarcinoma cell lines MDA-MB-231 were maintained in DMEM supplemented with 10% (v/v) fetal bovine serum (FBS) at 37°C in a humidified atmosphere containing 5% CO. Cells were routinely passaged using trypsin-EDTA following a wash with sterile PBS.

### Lentiviral production and generation of stable cell lines

HEK-293T cells were transfected using PolyJet transfection reagent (FroggaBio), according to the manufacturer’s instructions. The lentiviral packaging system consisted of the plasmid of interest (2 µg), HDM-VSVG (0.15 µg), HDM-tat1b (0.075 µg), HDM-HgPM2 (0.075 µg) and pRC-CMV-Ral1 (0.075 µg) per 10 cm^2^ well. At 18 h post-transfection, the culture medium was replaced with fresh medium. Viral supernatants were collected 48–72 h after transfection, filtered through 0.45-µm filters and stored at −80°C until use.

MDA-MB-231 cells were transduced with different volumes of viral supernatants for 18–24 h. Following medium replacement, puromycin selection was initiated at a concentration of 1 µg/mL and maintained thereafter. Selection was continued until death of all non-infected control cells. Surviving cells were expanded and cryopreserved in freezing medium containing 50% FBS, 40% DMEM and 10% DMSO.

### Protein extraction and western blot

Cells were washed with cold PBS, scraped and pelleted by centrifugation. Cell pellets were lysed in cold RIPA buffer (1% triton x100, 1% NaDOC, 1mM EDTA pH 8,0, 50mM Tris HCl, 0,1% SDS, 1% NP40, 50mM NaCl) supplemented with freshly added protease inhibitors. Lysates were vortexed three times for 10 s, with 5-min intervals on ice, and incubated for 1 h at 4°C with agitation. Samples were subsequently sonicated twice for 3 s at 12% intensity, with a 5-min interval on ice, and centrifuged at 13,000 rpm for 20 min. Supernatants were collected for protein quantification. Protein concentrations were determined using the Pierce BCA Protein Assay (ThermoFisher Scientific), according to the manufacturer’s instructions. Absorbance was measured at 562 nm using a microplate reader. Equal amounts of protein were prepared for each experimental condition, mixed with 4× Laemmli buffer (60 mM Tris-HCl pH 6.8, 2% SDS, 10% glycerol, 5% 2-mercaptoethanol, and 0.5% bromophenol blue) and heated at 98°C for 6 min.

Protein samples (15 µg per lane) were separated by SDS-PAGE using 8–15% resolving gels, depending on the molecular weight of the target proteins, with a 4.8% stacking gel. Electrophoresis was performed at 86 V for 20 min, followed by 116 V for approximately 90 min in running buffer containing 25 mM Tris base, 200 mM glycine and 0.1% SDS. Proteins were transferred onto nitrocellulose membranes (0.22 or 0.45 µm, depending on protein size) using cold transfer buffer containing 25 mM Tris base, 200 mM glycine and 20% ethanol. Transfer was performed at 116 V for 90 min at room temperature. Membranes were blocked for 30 min in 5% milk prepared in TBS-T (20 mM Tris-HCl, 150 mM NaCl and 0.1% Tween-20) and incubated overnight at 4°C with primary antibodies diluted in blocking buffer. Following three 5-min washes with TBS-T, membranes were incubated with HRP-conjugated secondary antibodies for 2 h at room temperature. Membranes were washed three times with TBS-T and once with TBS before chemiluminescent detection. Strong signals were detected using an in-house ECL substrate containing luminol and coumaric acid. For weaker signals, Clarity Western ECL or Clarity Max Western ECL substrates (Bio-Rad) were used. Images were acquired using an iBright FL1500 imaging system (ThermoFisher Scientific), and band intensities were quantified using Fiji software (version 2.16.0).

For sequential antibody detection, membranes were stripped and reprobed as required. When primary antibodies originated from different species, membranes were incubated in 0.01% acetic acid for 10 min at room temperature, followed by two washes with TBS-T. For antibodies from the same species, mild stripping was performed using buffer containing 200 mM glycine, 0.1% SDS and 1% Tween-20 (pH 2.5) for 10 min at room temperature. When stronger stripping was required, membranes were incubated for 10 min at 50°C in buffer containing 62.5 mM Tris-HCl (pH 6.8), 1% SDS and 0.8% β-mercaptoethanol. Membranes were subsequently washed and reblocked before incubation with the next primary antibody.

### MTT cell viability assay

The MTT stock solution (BioBasic) was prepared at 50 mg/mL in sterile PBS, stirred for 2 h at room temperature protected from light, filtered through a 0.22-µm filter and stored at −20°C in the dark. Fresh stock solutions were prepared at least every four months. On the day of the experiment, the stock was diluted in complete DMEM containing 10% FBS to obtain a 5 mg/mL working solution. MDA-MB-231 cells were seeded in 96-well plates at 10,000 cells per well. After 24 h, cells were treated with different concentrations of 5-fluorouracil (5-FU), doxorubicin (DOX) or cisplatin (CIS). Following 12 h of treatment, different concentrations of recombinant TRAIL were added, and cells were incubated for an additional 12 h. All treatments were performed in complete DMEM. Following treatment, 10 µL of MTT working solution (5 mg/mL) was added to each well to obtain a final concentration of 0.5 mg/mL. Cells were incubated for 4 h at 37°C in a humidified atmosphere containing 5% CO. The culture medium was then removed, and 100 µL of prewarmed DMSO was added to each well to dissolve the formazan crystals. Plates were incubated for 10 min at 37°C and subsequently agitated at 300–500 rpm for 20 min at room temperature. Absorbance was measured at 570 nm using an Epoch 2 Microplate Spectrophotometer reader (BioTek).

### Statistical analysis

Statistical analyses were performed using GraphPad Prism (version 9.3.1). Data normality was assessed using the Shapiro–Wilk test. Normally distributed data were analyzed using one-way analysis of variance (ANOVA), whereas the Kruskal–Wallis test was used for non-normally distributed data. The effects of chemotherapy and TRAIL treatment on cell viability, including potential interactions between these two factors, were assessed using two-way ANOVA followed by Dunnett’s multiple comparisons test. Statistical significance was set at p < 0.05, with significance levels indicated as follows: p < 0.05 (*), p < 0.005 (**), p < 0.001 (***) and p < 0.0001 (****). Non-significant differences are indicated by the absence of a symbol.

## Results

### Structural impact of exon 3-skipping on TRAIL-R4

TRAIL-R4 canonical transcript comprises nine exons and encodes the full-length TRAIL-R4 receptor, which contains an extracellular region composed of three cysteine-rich domains (CRD), a transmembrane domain (TM), and a truncated intracellular death domain (DD). The extracellular region also harbors the pre-ligand assembly domain (PLAD), located within CRD1, which contributes to receptor oligomerization and organization at the plasma membrane. Alternative splicing involving exon 3 skipping generates a shorter transcript in which exon 2 is directly joined to exon 4, resulting in the loss of the exon 3-encoded sequence. This event removes a substantial portion of the extracellular region, affecting both CRD1 and CRD2 and potentially altering the structural integrity of the PLAD-containing region (**Figure 1A**). Translation of this transcript produces a truncated TRAIL-R4 isoform, named iTRAIL-R4, that retains portion of CRD1 and CRD3 (**Figure 1B**).

**Figure 1.**
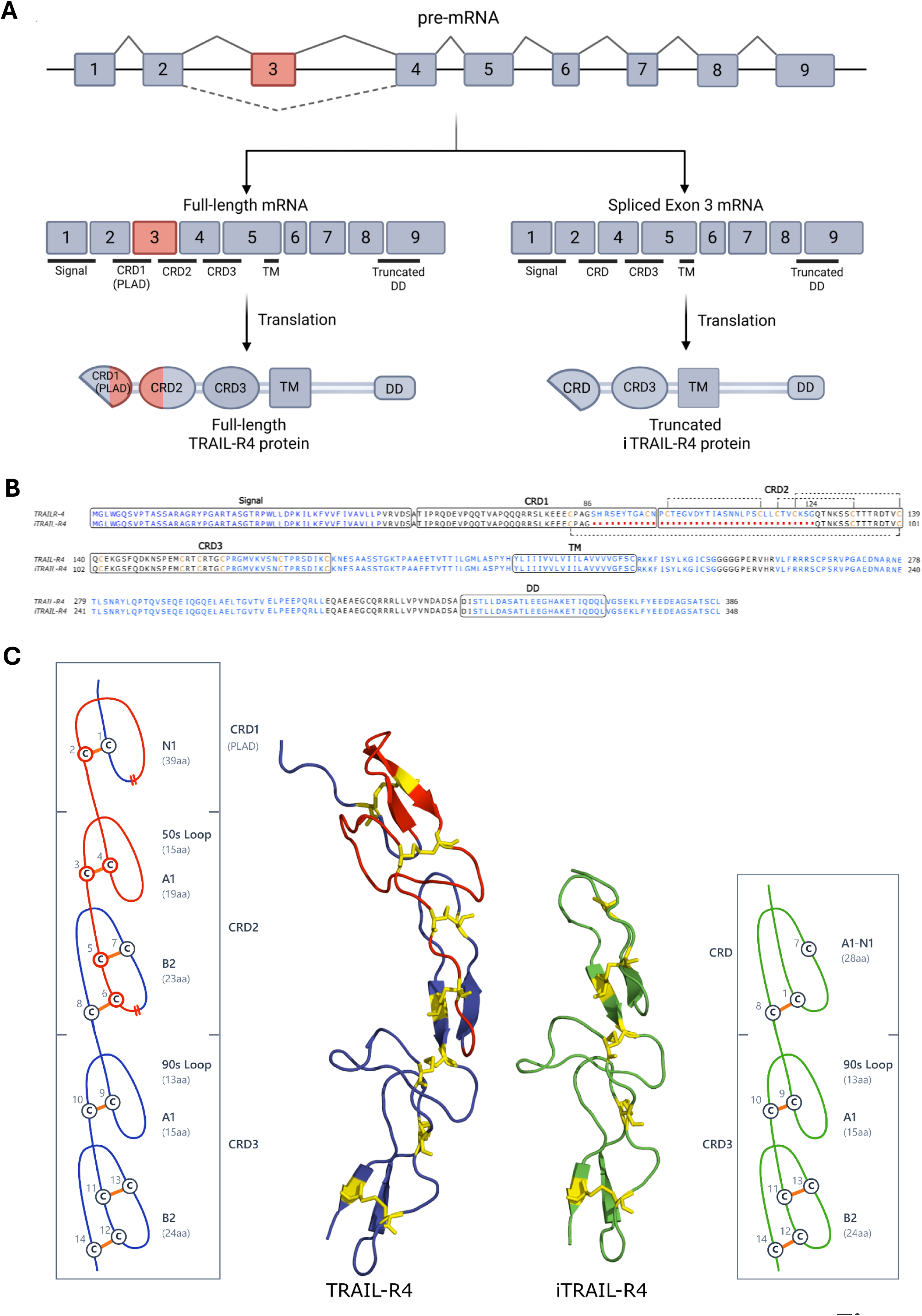
Alternative splicing of TRAIL-R4 and structural comparison of its isoforms. **(A)** Schematic representation of the TRAIL-R4 pre-mRNA, comprising nine exons, and the alternative splicing event resulting in exon 3 skipping (highlighted in red). The resulting full-length and exon 3-skipped mRNA transcripts are shown, along with their corresponding protein domains: signal peptide, cysteine-rich domains (CRD1/PLAD, CRD2, and CRD3), transmembrane domain (TM), and truncated death domain (DD). The bottom panels illustrate the resulting protein isoforms: full-length TRAIL-R4 and the truncated iTRAIL-R4 isoform. **(B)** Protein sequence alignment of full-length TRAIL-R4 and iTRAIL-R4. Cysteine residues involved in disulfide bond formation are highlighted in red. **(C)** Schematic two-and three-dimensional ribbon representations of the extracellular domains of full-length TRAIL-R4 (left) and truncated iTRAIL-R4 (right), showing CRD1 (N1), CRD2 (A1 and B2), and CRD3 (A1 and B2). Disulfide bonds are highlighted in yellow, and the region encoded by exon 3 is highlighted in red.

TNF receptor superfamily members share a conserved extracellular ligand-binding architecture composed of cysteine-rich domains (CRDs) (Locksley et al., 2001) (**Supplementary figure 1A**). TRAIL receptors contain three CRDs and adopt an elongated conformation stabilized by seven disulfide bonds (Chan et al., 2000). In the TRAIL-R4 protein, the first disulfide bond connects cysteine residues 83 and 96 within CRD1 (residues 57-97), while the remaining disulfide bonds six are equally distributed between CRD2 (residues 98-139) and CRD3 (residues 140-180) (**Figure 1B**). Each CRD can be subdivided into distinct structural modules. CRD1 contains the N1 module while CRD2 and CRD3 each comprise two modules, designated as A1 and B2. The A1 module contains one disulfide bond (Cys1–Cys2), whereas B2 contains two alternating disulfide bonds (Cys1–Cys3 and Cys2–Cys4) (**Figure 1C**). The A1 modules of CRD2 and CRD3 form the 50s and 90s loops, respectively, named according to their amino acid positions following signal peptide cleavage. These loops are primarily involved in ligand interactions (Cha et al., 1999).

TRAIL-R4 splicing event affects both CRD1 and CRD2 domains. We showed that exon 3 skipping substantially remodels the extracellular structure of TRAIL-R4 receptor, potentially affecting the integrity of the CRD1/PLAD domain and its ability to mediate receptor oligomerization. Although the transmembrane and intracellular regions remain intact, alterations to the extracellular architecture may influence receptor assembly, ligand binding and downstream ligand-mediated signalling.

### Consequences of TRAIL-R4 alternative splicing on ligand binding

Beyond their individual structural organization, TRAIL-R4 receptors can interact with one another through extracellular receptor-receptor interactions, promoting the formation of higher-order oligomeric assemblies at the membrane (**Supplementary figure 1B**). These interactions are mediated, in part, by the PLAD domain which facilitates receptor association independently of ligand binding. Following ligand binding, the formation of multimeric receptor complexes further enhances receptor clustering at the membrane (**Supplementary figure 1C**). Structural alterations affecting the PLAD-containing region, such as those resulting from exon 3 skipping in TRAIL-R4, could interfere with receptor-receptor interactions and alter the organization of receptor honeycomb complexes at the membrane.

To investigate the consequence of TRAIL-R4 splicing on the ligand binding to the receptor, we performed a detailed structural analysis of the contacts between the TRAIL-R4 receptor and its TRAIL ligand. We generated structural models using AlphaFold and aligned in PyMOL to the TRAIL:TRAIL-R2 complex (PDB: 1D0G). Our results confirmed that canonical TRAIL-R4 contained two negatively charged zones corresponding respectively to the receptor 90s loop located in CRD3 (zone A), and 50s loop located in CRD2 (zone B) (**Figure 2B**). Zone A, which primarily contributes to ligand specificity and cross-reactivity, is defined by contacts between the receptor 90s loop and Gln-205 of TRAIL. Gln-205 of TRAIL directly interacts with Glu-153 and Met-154 of TRAIL-R4. Other TRAIL residues, including Asn-199, Lys-201, Asp-203, and Glu-236, also participate in polar interactions, notably with Arg-156, Arg-104, and Asp-177 of TRAIL-R4 (**Figure 2B, Supplementary figure 2A**). Zone B forms a characteristic hydrophobic interface and contributes substantially to ligand–receptor interaction stability. This region forms structural contacts with the TRAIL α and β subunits. The interaction, between Tyr-216 in TRAIL and Leu-116 of TRAIL-R4, forms the core of a hydrophobic cluster further stabilized by the side chains of Ala-108, Asn-110 and Leu-112 (**Supplementary figure 2B**).

**Figure 2.**
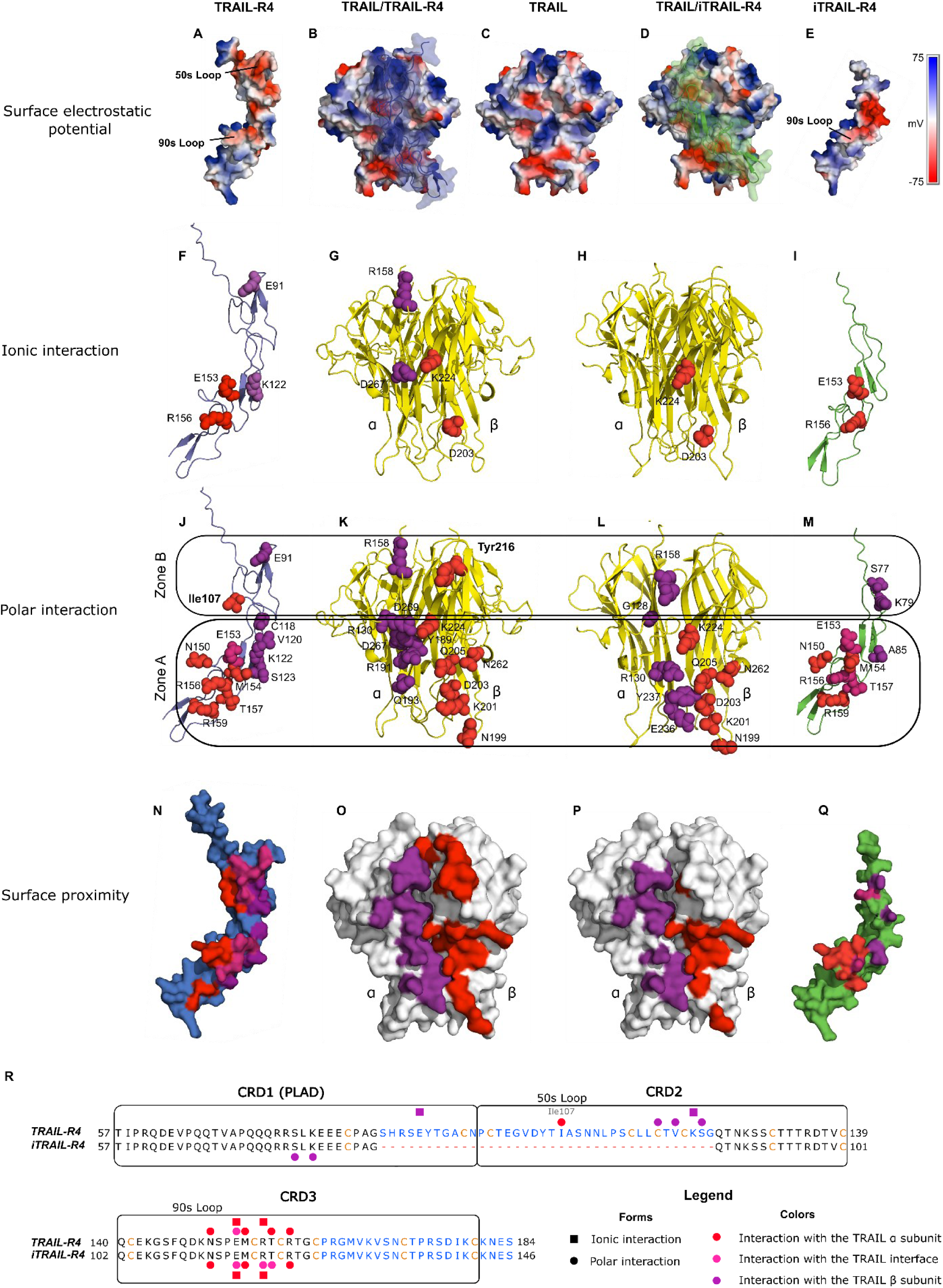
Structural characterization of TRAIL–receptor interactions. (A–E) Electrostatic surface potentials of TRAIL-R4, TRAIL:TRAIL-R4 complex, TRAIL, TRAIL:isoTRAIL-R4 complex, and isoTRAIL-R4. Red and blue indicate negative and positive charges, respectively. **(F–I)** Charged residues involved in ionic interactions mapped by PyMOL on TRAIL-R4, TRAIL (in complex with TRAIL-R4 or isoTRAIL-R4), and isoTRAIL-R4. **(J–M)** Residues involved in polar interactions between TRAIL and TRAIL-R4, and between TRAIL and isoTRAIL-R4. **(N–Q)** Contact surfaces between TRAIL-R4 and TRAIL, and TRAIL and isoTRAIL-R4. **(R)** Sequence alignment of canonical and isoform TRAIL-R4 highlighting residues interacting with TRAIL. Symbols denote ionic and polar interactions predicted by AlphaFold Multimer. Color code: TRAIL-R4 (blue), TRAIL (yellow and white), iTRAIL-R4 (green); α subunit interactions (purple), β subunit (red), interface (dark pink).

Our analysis also showed that iTRAIL-R4 isoform lacked the 50s loop and instead displayed an extended electronegative surface (**Figure 2E**). However, the 90s loop remains conserved. The iTRAIL-R4 receptor establishes contacts with both α and β subunits of TRAIL. In the canonical TRAIL-R4 form, multiple ionic interactions were observed, including Glu91–Arg158 (2.6 Å) and Lys122–Asp67 (3.4 Å) within the α subunit, as well as Glu153–Lys224 (3.6 Å) and Arg156–Asp203 (4.8 Å) within the β subunit (**Supplemental figure 2A**). In the iTRAIL-R4 isoform, interactions involving the α subunit (Glu91 and Lys122) were lost, whereas β subunit interactions were largely preserved (**Figure 2I**). Polar interaction analysis confirmed a reduction in contacts with the α subunit, partially compensated by the emergence of new interactions, although a net loss is observed (**Figure 2M**). Only the Glu153–Arg191 interaction at the α/β interface was conserved (**Supplementary figure 2C**). Most β subunit interactions remained intact however, the critical Tyr216 interaction, essential for TNF family receptor binding, was abolished. Notably, all the residues contributing to contact zone B were absent in iTRAIL-R4. Surface analysis further revealed a reduction in TRAIL-contacting zone B (**Figure 2Q**). These findings were consistent with AlphaFold-Multimer predictions of the TRAIL:iTRAIL-R4 complex, which show no restoration of interactions in this region.

Collectively, these structural analyses indicated that exon 3 skipping profoundly alters key determinants of ligand recognition. Although iTRAIL-R4 retains some interactions with TRAIL, the loss of critical residues within the 50s loop and the disruption of several polar and hydrophobic contacts substantially modified the predicted receptor–ligand interface. These findings provide a structural rationale for the altered TRAIL-binding properties of iTRAIL-R4 and suggest that alternative splicing of TRAIL-R4 influence both ligand recognition and receptor organization.

### iTRAIL-R4 exhibits reduced anti-apoptotic activity

TRAIL-R4 is expressed at relatively low levels in MDA-MB-231 breast cancer cells, which exhibit concentration-dependent sensitivity to TRAIL (Avi et al., 1999; Avi & m., 1999; Falschlehner et al., 2007; Pitti et al., 1996; Sanlioglu et al., 2005). Their pronounced sensitivity to TRAIL, combined with low endogenous TRAIL-R4 expression, provides a suitable experimental context to determine whether TRAIL-R4 and iTRAIL-R4 expression modulate the cellular response to TRAIL. To this end, MDA-MB-231 cells were transduced with lentiviral vectors encoding either TRAIL-R4 or iTRAIL-R4, or with a control vector, followed by puromycin selection to establish stable cell lines. The resulting cell lines were treated with TRAIL (200 ng/mL, 6 h), and activation of the canonical apoptotic signaling cascade downstream of TRAIL receptor engagement, as well as markers of mitochondrial apoptotic pathway activation, were assessed by Western blot (**Figure 3A**). TRAIL-R4 overexpression in MDA-MB-231 cells strongly inhibited apoptosis, as indicated by decreased levels of caspase-9 (-69%), caspase-8 (-82.5% and - 60%), caspase-7 (-75%), caspase-3 (-68%), PARP (-68%), γH2AX (-67%), tBid (-76%), and RIP (-45%), corresponding to an average reduction of ∼68.5% (****p < 0.0001) (**Figure 3B**). In contrast, isoform-expressing cells exhibited higher apoptotic activity compared to TRAIL-R4-expressing cells, with modest changes relative to wild-type (WT) cells. Minor decrease in caspase-9 (–28%, ***) and caspase-3 (–31%, **) were observed but no significant changes for other markers (**Figure 3B**). The overall decrease in apoptotic signaling (∼11%, **p < 0.01) indicated a substantially weaker inhibitory effect. To determine whether the reduced apoptotic signaling observed upon canonical TRAIL-R4 overexpression was associated with enhanced anti-apoptotic signaling, we examined the expression of key regulators of apoptosis. Consistent with the reduced caspase activation, XIAP and Bcl-xL protein levels were higher in cells expressing canonical TRAIL-R4 than in WT or iTRAIL-R4-expressing cells. Conversely, cleaved RIP levels were elevated in WT and iTRAIL-R4-expressing cells, further supporting the enhanced apoptotic response observed in these conditions. In contrast, TRADD levels remained unchanged across conditions, suggesting that the differential apoptotic response was not associated with changes in TRADD expression. Collectively, these findings demonstrate that iTRAIL-R4 lacks the potent anti-apoptotic activity of canonical TRAIL-R4. These results identify alternative splicing of TRAIL-R4 as a key determinant of its anti-apoptotic function and suggest that shifting the balance toward iTRAIL-R4 could provide a promising strategy to sensitize cancer cells to TRAIL-induced apoptosis.

**Figure 3.**
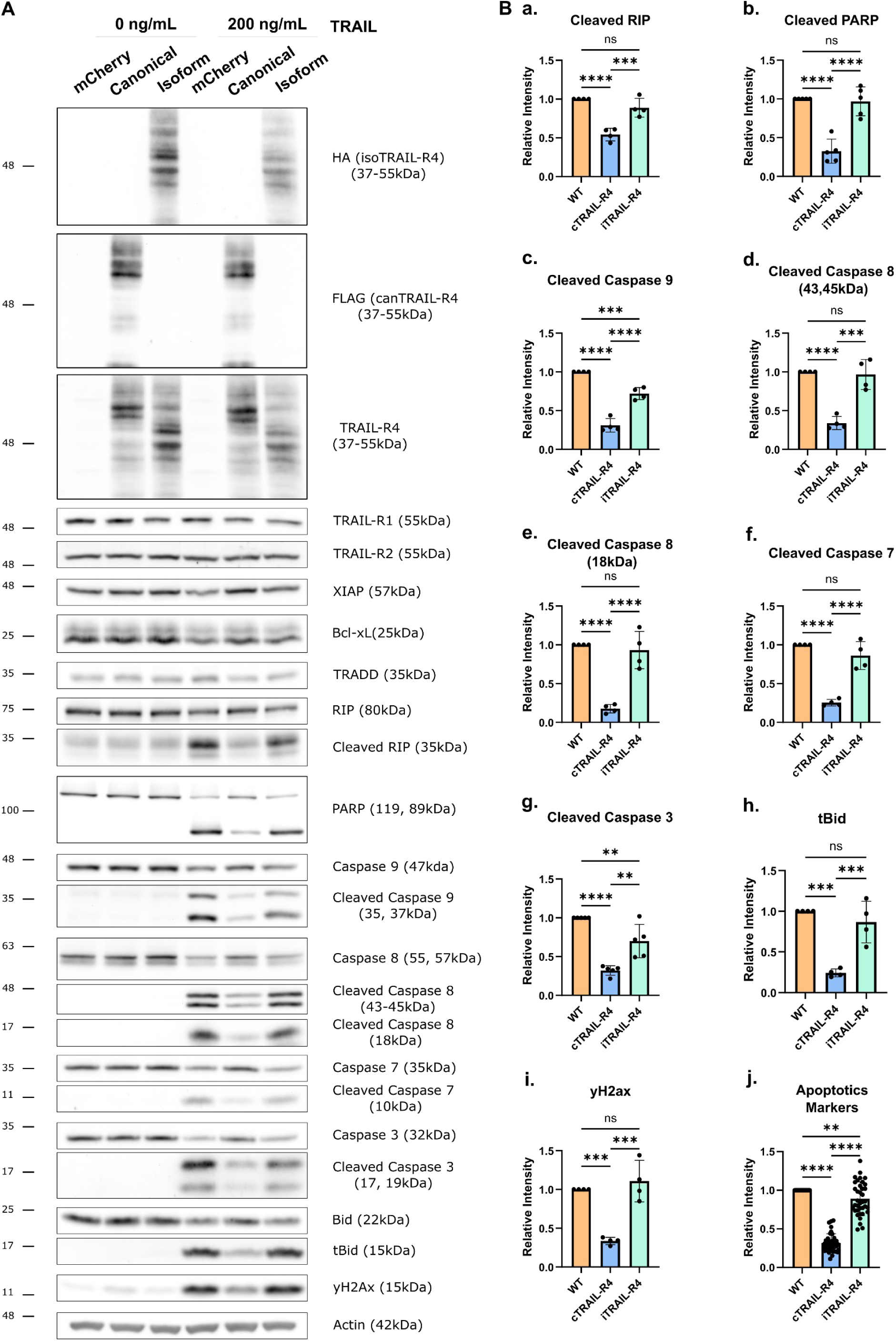
TRAIL-mediated apoptosis in MDA-MB-231 cells. **(A)** MDA-MB-231 cells were treated with TRAIL (200 ng/mL, 6 h) and analyzed by Western blot for TRAIL-R1/DR4 and TRAIL-R2/DR5 expression, extrinsic (caspases-8,-3,-7) and intrinsic (BID, caspase-9) apoptosis markers, general apoptosis markers (PARP, γH2AX), anti-apoptotic proteins (XIAP, Bcl-xL), and non-apoptotic TRAIL signaling components (TRADD, RIP). **(B)** Quantification of immunoblots. Signals were normalized to actin and expressed relative to wild-type cells. Data are presented as mean ± SD (n = 3-4); statistical analysis was performed using ANOVA with multiple comparisons (*p < 0.05, **p < 0.01, ***p < 0.001, ****p < 0.0001).

### iTRAIL-R4 antagonizes the anti-apoptotic activity of TRAIL-R4

To determine whether the effects of TRAIL-R4 overexpression were associated with changes in TRAIL receptor expression, we examined TRAIL-R1 and TRAIL-R2 protein levels. TRAIL-R1 expression was reduced by approximately 33% upon overexpression of either TRAIL-R4 or iTRAIL-R4 (**Figure 4A**; p < 0.05), whereas TRAIL-R2 expression remained unchanged (**Figure 4B**). This selective reduction in TRAIL-R1 levels may be relevant in light of previous studies demonstrating that TRAIL-R1, TRAIL-R2, and TRAIL-R4 can form both homodimers and ligand-independent heterodimers (Neumann et al., 2014). In particular, the ability of TRAIL-R4 to heterodimerize with TRAIL-R1 and TRAIL-R2 has led to the hypothesis that TRAIL-R4 may antagonize death receptor signaling not only by competing for ligand binding, but also by incorporating into receptor complexes and interfering with the receptor organization required for efficient apoptotic signaling (Mérino et al., 2006; Reis et al., 2010; Szegezdi, Sloot, et al., 2012; Wajant, 2019b). Our observation that canonical or isoform TRAIL-R4 overexpression selectively reduced TRAIL-R1 levels may therefore suggest a preferential relationship between these two receptors.

**Figure 4.**
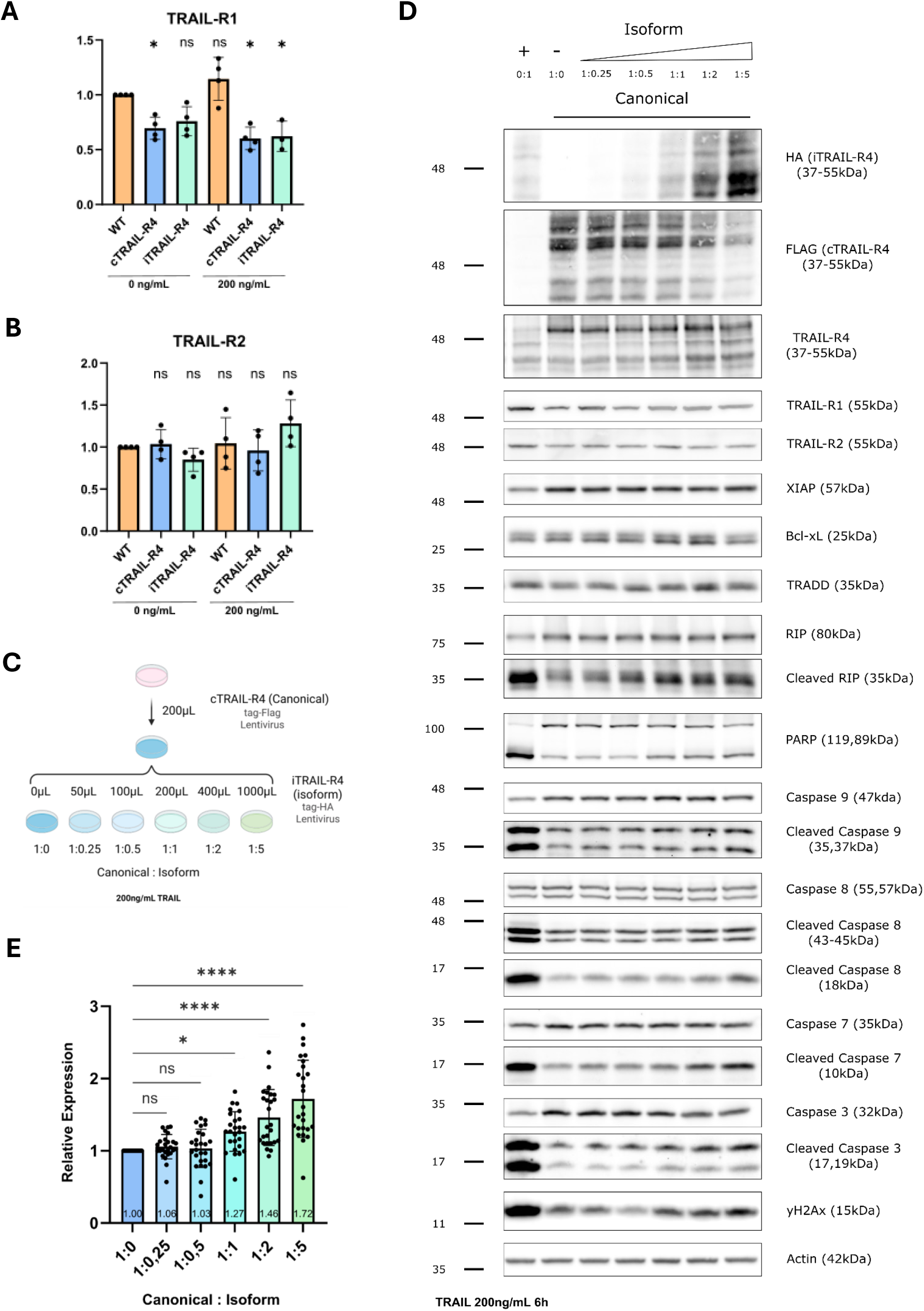
TRAIL-R4 isoform ratio in regulating TRAIL-induced apoptosis. (A-B) Cells were treated with TRAIL (200 ng/mL, 6 h) and analyzed by Western blot for TRAIL-R1 **(A)** and TRAIL-R2 **(B)**. Signals were quantified, normalized to actin, and expressed relative to untreated WT. Data are mean (n = 3–4); ANOVA with multiple comparisons (*p < 0.05, **p < 0.01, ***p < 0.001, ****p < 0.0001). **(C)** Schematic representation of the experimental design. **(D)** Cells were treated with TRAIL (200 ng/mL, 6 h) and analyzed by Western blot for TRAIL-R1 and TRAIL-R2 expression, extrinsic (caspases-8,-3,-7) and intrinsic (BID, caspase-9) apoptosis markers, general apoptosis markers (PARP, γH2AX), anti-apoptotic proteins (XIAP, Bcl-xL), and non-apoptotic TRAIL signaling components (TRADD, RIP). **(E)** Quantification of immunoblots. Signals were normalized to actin and expressed relative to WT. Data are presented as mean ± SD (n = 3–4); statistical analysis was performed using ANOVA with multiple comparisons (*p < 0.05, **p < 0.01, ***p < 0.001, ****p < 0.0001).

We next asked whether the iTRAIL-R4 expression could counteract the anti-apoptotic activity of canonical TRAIL-R4. Cells stably overexpressing TRAIL-R4 were transduced with increasing levels of iTRAIL-R4, generating canonical-to-isoform expression ratios ranging from 1:0 to 1:5 (**Figure 4C**). Reciprocal expression of the two isoforms was confirmed by detection of HA-tagged iTRAIL-R4 and Flag-tagged canonical TRAIL-R4. Consistent with our previous findings, TRAIL-R1 expression was reduced under TRAIL-R4-overexpressing conditions, whereas TRAIL-R2 levels remained unchanged. As expected, overexpression of TRAIL-R4 markedly suppressed apoptotic signaling compared with control cells (1:1). Strikingly, increasing iTRAIL-R4 expression progressively reversed this anti-apoptotic effect, restoring apoptotic signaling by 5% (1:0.25), 3% (1:0.5), 26% (1:1), 46% (1:2), and 72% (1:5) (**Figure 4D-E**). This reversal became statistically significant at canonical-to-isoform ratios of 1:1 and higher, demonstrating a clear dose-dependent relationship between iTRAIL-R4 expression and the restoration of apoptotic signaling. Thus, increasing iTRAIL-R4 expression progressively counteracts the protective effect conferred by canonical TRAIL-R4. Collectively, these results provide strong evidence that iTRAIL-R4 functionally antagonizes the anti-apoptotic activity of canonical TRAIL-R4. Importantly, the progressive restoration of apoptotic signaling as the relative abundance of iTRAIL-R4 increases suggests that the balance between the two isoforms is a critical determinant of cellular sensitivity to TRAIL.

### iTRAIL-R4 sensitizes cancer cells to TRAIL-based combination therapies

The ability of TRAIL to enhance the cytotoxic effects of chemotherapeutic agents has been widely documented, particularly in combination with 5-fluorouracil (5-FU) (Avi et al., 1999; Li et al., 2021; Nazim et al., 2017; Zhao et al., 2012; Zhu et al., 2013), doxorubicin (Wang et al., 2010), and cisplatin (Woods et al., 2008; Yin et al., 2011). Building on these findings, we investigated whether the pro-apoptotic effect associated with iTRAIL-R4 expression could be further enhanced by combining TRAIL with conventional chemotherapeutic agents. Cell viability was assessed by MTT assays following treatment with TRAIL alone or in combination with doxorubicin, cisplatin, or 5-FU (**Figure 5**). As previously observed, canonical TRAIL-R4 overexpression conferred st substantial resistance to TRAIL-induced cytotoxicity, resulting in approximately 75% cell viability at 1,000 ng/mL TRAIL, compared with 52% in iTRAIL-R4-expressing cells and 28% in control cells (**Figure 5A-a**). Notably, the addition of doxorubicin markedly enhanced TRAIL-mediated cytotoxicity, shifting the dose-response curves toward reduced cell viability, particularly in control and iTRAIL-R4-expressing cells (**Figure 5A-b**). Importantly, treatment with doxorubicin alone (1 µM) resulted in relatively high cell viability (∼77%) in all cell lines, indicating that the enhanced cytotoxic response primarily resulted from the combination of doxorubicin and TRAIL rather than from doxorubicin treatment alone (**Figure 5A-c**). Cisplatin produced a similar, although less pronounced, enhancement of TRAIL-induced cytotoxicity (**Figure 5B**). In contrast, 5-FU had a limited effect following a 24h treatment (**Supplementary figure 3**). Collectively, these findings demonstrate that iTRAIL-R4 expression enhances cancer cell susceptibility to TRAIL-based combination treatments, particularly those involving doxorubicin and cisplatin. In contrast, cells overexpressing canonical TRAIL-R4 remained substantially more resistant to these treatments. These results highlight the potential of iTRAIL-R4 to overcome TRAIL-R4-mediated resistance and improve the efficacy of TRAIL-based therapeutic combinations. They further support the therapeutic relevance of modulating TRAIL-R4 alternative splicing to shift cancer cells toward a more apoptosis-sensitive state.

**Figure 5.**
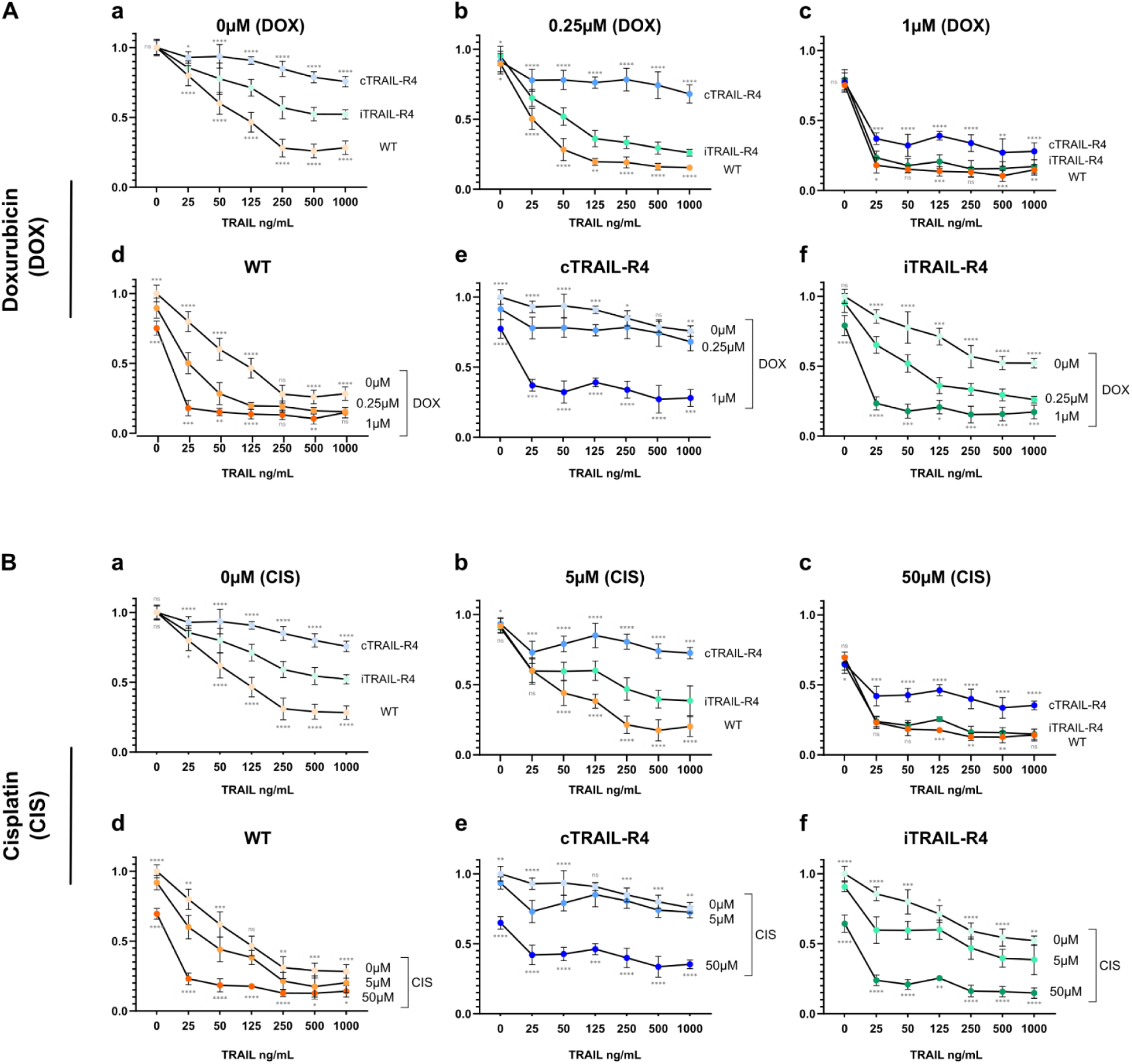
Chemotherapeutic sensitization to TRAIL-induced cytotoxicity. MDA-MB-231 cells, either wild-type (WT) or expressing canonical TRAIL-R4 (cTRAIL-R4) or iTRAIL-R4, were treated for 24 h with two concentrations of chemotherapeutic agents, with increasing concentrations of TRAIL (0–1000 ng/mL) added during the final 12 h. Cell viability was assessed by MTT assay and expressed relative to untreated cells. Responses to TRAIL in combination with **(A)** doxorubicin or **(B)** cisplatin are presented in panels a-c, organized by chemotherapeutic agent concentration, and d-f, organized by TRAIL-R4 expression status (WT, canTRAIL-R4, and isoTRAIL-R4). Data represent mean ± SD of three independent technical replicates. Statistical analyses were performed using two-way ANOVA with multiple comparisons against iTRAIL-R4-expressing cells (*p < 0.05, **p < 0.01, ***p < 0.001, ****p < 0.0001).

## Discussion

TRAIL-based therapies have attracted considerable interest because of their ability to preferentially induce apoptosis in cancer cells while largely sparing normal tissues. However, their clinical efficacy has been limited by intrinsic and acquired resistance mechanisms. Among these, TRAIL-R4 is an important regulator of cellular sensitivity to TRAIL. Lacking a functional death domain, TRAIL-R4 can suppress apoptosis through ligand sequestration, interference with receptor organization, and activation of pro-survival pathways. Here, we identified alternative splicing as an additional mechanism regulating TRAIL-R4 function. We demonstrated that iTRAIL-R4 has reduced anti-apoptotic activity compared with canonical TRAIL-R4 and progressively counteracts its protective effect as its relative expression increases. These findings reveal that TRAIL-R4 activity depends not only on receptor abundance but also on the balance between its distinct isoforms.

### iTRAIL-R4 structure

Our structural analyses provide a potential explanation for the functional differences between the two TRAIL-R4 forms. Alternative splicing resulting in exon 3 skipping removes part of the CRD2 domain and eliminates several residues predicted to contribute to TRAIL binding. In particular, the deletion disrupts the electrostatic and polar interaction network at the receptor–ligand interface. Among these interactions, the contact between Y216 of TRAIL and L116 of TRAIL-R4 is particularly notable. Importantly, L116 is absent from iTRAIL-R4, providing a structural rationale for the predicted reduction in ligand interactions. The functional importance of this region is further supported by previous mutagenesis studies showing that Y216 is a critical residue for bioactivity and ligand binding not only to TRAIL-R4 but also to other TNF receptor family members, including TNFα, TNFβ, and FasL (Goh & Porter, 1991; Hymowitz et al., 1999; ichi Yamagishi et al., 1990; Ostade et al., 1994; Schneider et al., 1997). But the potential consequences of exon 3 skipping event may extend beyond this interaction site. It disrupts multiple contacts across the receptor-ligand interface rather than eliminating a single critical interaction. Although structural modeling predicts the formation of new compensatory contacts in iTRAIL-R4, these appear insufficient to restore the interaction network observed in the canonical receptor.

Importantly, exon 3 skipping also affects part of the PLAD domain, raising the possibility that iTRAIL-R4 may exhibit altered receptor oligomerization. TRAIL receptors can form homo-and heterodimeric complexes independently of ligand binding, including interactions between TRAIL-R4 and the apoptotic receptors TRAIL-R1 and TRAIL-R2 (Lee et al., 2005b; Neumann et al., 2014). These interactions suggest that the anti-apoptotic activity of TRAIL-R4 cannot be explained solely by competition for ligand binding, but may also involve modulation of receptor organization at the plasma membrane (Mérino et al., 2006; Reis et al., 2010; Szegezdi, Sloot, et al., 2012; Wajant, 2019b). Consistent with this possibility, Clancy et al., 2005 demonstrated that a TRAIL-R4 mutant unable to bind TRAIL but retaining an intact PLAD could still protect cells against TRAIL-induced apoptosis, supporting a role for ligand-independent receptor interactions in its inhibitory function. In our model, although exon 3 skipping removes a substantial portion of the region corresponding to the canonical PLAD, structural predictions suggest that a disulfide bridge may still form in iTRAIL-R4. Whether this structure is sufficient to preserve interactions with other TRAIL receptors remains unknown. Thus, we cannot conclude that iTRAIL-R4 has completely lost its capacity for receptor association. Nevertheless, the extensive deletion is likely to alter the structural organization of this region and may consequently affect receptor assembly and its ability to interfere with apoptotic signaling. Collectively, our findings suggest that exon 3 skipping may affect TRAIL-R4 function through two complementary mechanisms: reduced ligand engagement and altered receptor organization. Both could contribute to the distinct functional properties of iTRAIL-R4.

### Functional consequences of TRAIL-R4 alternative splicing

Our functional data support the structural predictions, demonstrating that iTRAIL-R4 has substantially reduced anti-apoptotic activity compared with canonical TRAIL-R4. Whereas the canonical receptor strongly suppressed TRAIL-induced apoptosis, cells expressing iTRAIL-R4 retained greater caspase activation and apoptotic signaling. Notably, iTRAIL-R4 did not simply represent a loss-of-function variant, as increasing its expression progressively restored apoptotic signaling in cells overexpressing canonical TRAIL-R4. This finding suggests that iTRAIL-R4 can counteract the protective activity of the canonical receptor, potentially by competing for receptor assembly at the plasma membrane, forming hetero-oligomeric complexes with reduced inhibitory capacity, or interfering with receptor trafficking and surface localization. Although these mechanisms remain to be established, the progressive restoration of apoptosis indicates that the relative abundance of the two isoforms, rather than total TRAIL-R4 expression alone, may be an important determinant of cellular sensitivity to TRAIL. Alternative splicing may therefore provide a means of fine-tuning TRAIL-R4 activity by generating an isoform that retains some receptor properties while lacking the full anti-apoptotic capacity of the canonical protein. Beyond apoptosis regulation, TRAIL-R4 has also been implicated in non-apoptotic signaling pathways, including NF-κB, MAPK, and PI3K/AKT, which can influence cell survival, proliferation, and inflammatory responses depending on the cellular context (Avi et al., 1999; Avi & m., 1999; Degli-Esposti et al., 1997b; Hu et al., 1999; Liu et al., 2017; Ravi et al., 2001; Schneider et al., 1997). Given the ability of TRAIL-R4 to associate with apoptotic TRAIL receptors, alternative splicing could potentially influence these pathways by altering receptor organization and the recruitment of downstream signaling components. Whether iTRAIL-R4 participates in heteromeric receptor complexes, and whether such complexes differ in their capacity to activate non-apoptotic signaling, remains unknown. Investigating these pathways will therefore be important to determine whether TRAIL-R4 alternative splicing influences cellular responses to TRAIL beyond the regulation of apoptosis.

### TRAIL-R4 splicing and chemotherapy response

The therapeutic potential of iTRAIL-R4 must be considered in the broader context of strategies aimed at overcoming TRAIL resistance. Previous studies have demonstrated that combining TRAIL with conventional chemotherapeutic agents can restore apoptotic responses in otherwise resistant tumor cells (Belyanskaya et al., 2007; Li et al., 2021; Nazim et al., 2017; Ouyang et al., 2018; Wang et al., 2010; Woods et al., 2008; Xu et al., 2003). In particular, chemotherapy can partially overcome TRAIL-R4-mediated resistance, potentially by enhancing apoptotic signaling (Ganten et al., 2004). Our results highlight the importance of treatment sequence, as pretreatment with doxorubicin or cisplatin followed by TRAIL produced a marked increase in apoptotic sensitivity. This is consistent with previous evidence that chemotherapy can enhance TRAIL signaling by increasing caspase-8 recruitment and activation at the DISC, reducing c-FLIP expression, and increasing TRAIL-R2 levels (Ganten et al., 2004). The sensitizing effects of chemotherapy on TRAIL-induced apoptosis have been reported for several agents, including 5-fluorouracil (5-FU), doxorubicin, and cisplatin (Avi et al., 1999; Avi & m., 1999; Li et al., 2021; Nazim et al., 2017; Wang et al., 2010; Woods et al., 2008; Yin et al., 2011). Proposed mechanisms include increased expression of pro-apoptotic death receptors, such as TRAIL-R2 and Fas following 5-FU treatment (Wu & El-Deiry, 1997), as well as modulation of intracellular apoptotic regulators. Consistent with these findings, we observed stronger sensitization following doxorubicin and cisplatin treatment, whereas the response to 5-FU was more limited at 24 h and became more pronounced after 48 h. This difference suggests that the effectiveness of combination treatments may depend on the specific mechanisms and kinetics of action of each chemotherapeutic agent. Importantly, chemotherapy did not fully overcome the strong protective effect of canonical TRAIL-R4, particularly at lower concentrations. In contrast, cells expressing iTRAIL-R4 displayed substantially greater sensitivity following treatment, consistent with the reduced anti-apoptotic capacity of this isoform. These observations suggest that iTRAIL-R4 is less able than canonical TRAIL-R4 to maintain resistance when apoptotic signaling is potentiated by chemotherapy. Thus, alternative splicing may influence not only basal sensitivity to TRAIL but also the capacity of tumor cells to respond to combination therapies. Collectively, our findings identify the relative expression of TRAIL-R4 isoforms as a potential determinant of treatment response and suggest that modulating TRAIL-R4 alternative splicing could represent an additional strategy for improving the effectiveness of TRAIL-based anticancer therapies.

### Perspectives

Our findings expand the current understanding of TRAIL-R4-mediated resistance by identifying alternative splicing as an important regulator of receptor function. Whereas canonical TRAIL-R4 strongly protects cells against TRAIL-induced apoptosis, iTRAIL-R4 exhibits reduced anti-apoptotic activity and progressively counteracts the protective effect of the canonical receptor when co-expressed. Thus, the relative abundance of TRAIL-R4 isoforms may represent an additional determinant of cellular sensitivity to TRAIL and a potential target for therapeutic intervention. Several questions remain regarding the mechanisms underlying this functional antagonism. In particular, it remains to be determined whether iTRAIL-R4 competes with the canonical receptor for membrane localization or interferes with the formation and organization of TRAIL receptor complexes. Investigating potential interactions between iTRAIL-R4 and other TRAIL receptors using complementary biochemical and biophysical approaches will be essential to clarify its mode of action. Ultimately, determining whether TRAIL-R4 splicing can be therapeutically manipulated represents an important next step. Splice-switching strategies, including antisense oligonucleotides or small molecules, could potentially shift isoform expression toward iTRAIL-R4 and weaken TRAIL-R4-mediated resistance. Although further studies are needed to establish their feasibility and specificity, targeting the balance between TRAIL-R4 isoforms may offer a promising approach to sensitizing resistant tumor cells to TRAIL-based therapies.

## Acknowledgments

This effort was supported by a grant from the Canadian Institutes of Health Research to B.L. [PJT-166109]. C.P. was supported by a fellowship from the Université de Sherbrooke (Sherbrooke, Canada) and from the Canadian Institutes of Health Research.

