## Supplemental material for "A TRAIL-R4 splice isoform sensitizes cancer cells to TRAIL-induced apoptosis"

### Supplemental tables

### Supplemental table 1. Reagents and material

| Reagents and material | Supplier | Catalog number |
|---|---|---|
| **Chemicals and recombinant proteins** | | |
| Q5® High-Fidelity DNA Polymerase | New England BioLabs | #M0491 |
| NotI-HF | New England BioLabs | #R3189 |
| AscI | New England BioLabs | #R0558 |
| HindIII-HF | New England BioLabs | #R3104 |
| Antarctic phosphatase | New England BioLabs | #M0289 |
| T4 DNA Ligase | New England BioLabs | #M0202 |
| DMEM | Wisent | #319-005-CL |
| FBS | Avantor | #10803-034 |
| dPBS | Wisent | #311-425 CL |
| DMSO | Biobasic | #D0231 |
| NP-40 | ThermoFisher | #85124 |
| Sodium Deoxycholate | ThermoFisher | #89905 |
| P8340 Protease Inhibitor Cocktail | Millipore Sigma | #P8340 |
| Tween-20 | Millipore Sigma | #P1379 |
| Paraformaldehyde | ThermoFisher | #047340.9M |
| Normal Goat Serum | ThermoFisher | # PCN5000 |
| Triton X-100 | Millipore Sigma | # X100 |
| SlowFade Gold Antifade Mountant | ThermoFisher | #S36936 |
| Puromycin | BioShop | #PUR555 |
| Proteinase K | Biobasic | #PB0451 |
| Sodium Acetate | ThermoFisher | #AM9740 |
| EDTA | ThermoFisher | #15575020 |
| MTT (Thiazol blue tetrazolium bromide) | Biobasic | # T0793 |
| Acrylamide 30% | Biobasic | #A0011 |
| PolyJet | FroggaBio | #SL100688 |
| Puromycin | DSHB | #PMY-2A4-s |
| Trypsine/EDTA | Wisent | #325-043-EL |
| TRAIL/APO2L | GenScript USA inc | # Z03124-50 |
| WGA | AAT Bioquest | #25559(AAT) |
| DID | AAT Bioquest | #22034(AAT) |
| 5-Fluorouracil | ThermoFisher | #A13456.06 |
| Doxoribicin | ThermoFisher | #J64000.MA |
| Cisplatin | ThermoFisher | #010471.01 |
| **Kits** | | |
| Pierce BCA Protein Assay | ThermoFisher | #23225 |
| Clarity Western ECL substrate | Biorad | #1705061 |
| Clarity Max Western ECL substrate | Biorad | #1705062 |
| Gateway™ LR Clonase™ | ThermoFisher | #11791019 |

| | | |
|---|---|---|
| EZ-10 Spin Column Plasmid Miniprep Kit | BioBasic | # BS413 |
| EZ-10 Spin column PCR products purification | BioBasic | #BS363 |
| EZ-10 Spin column gel extraction purification | BioBasic | #9K-006-0002 |
| PureLink™ HiPure Plasmid Midiprep Kit | ThermoFisher | #K210004 |
| **Softwares et algorithms** | | |
| Fiji | (Schindelin et al., 2012) | https://fiji.sc/ |
| Prism (version 10.4.2) | GraphPad Software | https://www.graphpad.com/ |
| Biorender | Biorender | https://app.biorender.com |
| PyMOL | (Schrödinger, L. and DeLano, W. (2020) PyMOL.) | http://www.pymol.org/pymol |
| Alpha Fold | (Jumper et al., 2021) | Alphafold 3.0 multimer |
| **Data used in this study** | | |
| TRAIL-R1 | (Ramamurthy et al., 2015) | 5CIR |
| TRAIL-R2 | (Hymowitz et al., 1999) | 1D0G |
| | (Mongkolsapaya et al., 1999) | 1D4V |
| TRAIL-R4 | AlphaFold | AF-Q9UBN6-F1 |
| TNF | (Naismith et al., 1995) | 1NCF |
| Domaine transmembranaire TRAIL-R | (Pan et al., 2019) | 6NHY |
| **Antibodies** | | |
| β-Actin | Cell Signaling Technology | 4970 |
| α-Tubulin | Abclonal | A6830 |
| γH2AX | Cell Signaling Technology | 9718 |
| PARP | Cell Signaling Technology | 9542 |
| Caspase-3 | Abclonal | A19654 |
| Cleaved caspase-3 | Cell Signaling Technology | 9661 |
| Caspase-7 | Cell Signaling Technology | 9492 |
| Caspase-8 | Abclonal | A19549 |
| Caspase-8 | Abclonal | A0215 |
| Cleaved caspase-8 | Cell Signaling Technology | 9496 |
| Caspase-9 | Cell Signaling Technology | 9502 |
| BAX | Cell Signaling Technology | 2772 |
| Bcl-xL | Cell Signaling Technology | 2764 |
| cFLIP | Abclonal | A2555 |
| XIAP | Abclonal | A6869 |
| TRADD | Cell Signaling Technology | 3694 |
| RIP | Cell Signaling Technology | 3493 |
| Bid | Cell Signaling Technology | 2002 |
| TRAIL-R1/TNFRSF10A | Novus | NB100-56747SS |
| TRAIL-R2/TNFRSF10B | Novus | NB100-566 18SS |
| TRAIL-R4 | Abclonal | A6136 |
| Anti-HA | BioLegend | 901501 |
| Anti-FLAG | Cell Signaling Technology | 14793 |
| Anti-rabbit HRP | Jackson ImmunoResearch | 711-035-152 |
| Anti-mouse HRP | Jackson ImmunoResearch | 715-035-150 |

| Oligonucleotides | | Sequence |
|---|---|---|
| TRAIL-R4_NotI_F1 | Forward #1 | TCCGCGGCCGCCATGGGACTTT GGGGACAAAGCGTCC |
| TRAIL-R4_AscI_R1 | Reverse #1 | GTCGGCGCGCCCCAGGCAGGA CGTAGCAGAGCCTGC |
| iTRAIL-R4_HindIII_F2 | Forward #2 | AGGAAGCTTCCAGGATAAAAAC TCCCCTG |
| iTRAIL-R4_HindIII_R2 | Reverse #2 | TGGAAGCTTCCTTTTTCACACTG ACACACG |

**Supplementary figure legends**

**Supplementary figure 1. Structural organization and oligomerization of TRAIL receptors.** **(A)** Schematic representation of the four TRAIL receptors (TRAIL-R1, TRAIL-R2, TRAIL-R3, and TRAIL-R4), showing their extracellular cysteine-rich domains (CRD1–CRD3), transmembrane regions, and intracellular domains. CRD1 corresponds to the pre-ligand assembly domain (PLAD), which contributes to ligand-independent receptor association. TRAIL-R1 and TRAIL-R2 contain intracellular apoptotic (death) domains, whereas TRAIL-R3 lacks a functional apoptotic domain and TRAIL-R4 possesses a truncated apoptotic domain. **(B)** Schematic representation of the higher-order organization of TRAIL receptors, illustrating their potential assembly into an interconnected receptor network through extracellular interactions. **(C)** Proposed model of ligand-induced TRAIL receptor clustering. Binding of trimeric TRAIL ligand (yellow) promotes receptor association and oligomerization at the membrane, facilitating the recruitment of additional receptors and the formation of higher-order receptor complexes. This clustering promotes intracellular signaling through the receptor-associated apoptotic domains (red).

**Supplementary figure 2. Structural characterization of TRAIL–receptor interactions.** **(A)** Comparison of ionic and polar interactions between TRAIL-R4 and the TRAIL cytokine. Types of interactions identified between TRAIL residues (classified by α and β subunits as well as contact regions A and B) and residues of TRAIL-R4 receptor. Distances in Å are indicated when an interaction is present. An asterisk (*) marks residues involved in interactions with both TRAIL subunits (interface). Residues G128, R158, and R191 are part of the AA" loop of the α subunit of TRAIL. Interactions were identified from structural models visualized using PyMOL. **(B)** On the left panel, close-up showing a potential hydrogen bond between Tyr216 and Ile116 (3.0 Å). **(C)** Structural representations of the (a) TRAIL:iTRAIL-R4 complex, ionic interactions between (b) TRAIL and (c) iTRAIL-R4, and polar interactions between (d) TRAIL and (e) iTRAIL-R4. TRAIL is shown in yellow and iTRAIL-R4 in green. Interactions involving the α subunit of TRAIL are shown in purple, those involving the β subunit in red, and interactions at the interface between the two subunits in dark pink. **(D)** Comparison of ionic and polar interactions between iTRAIL-R4 and the TRAIL cytokine. A dash (–) indicates that no interaction was detected at that position. Colors highlight the following: Red: interactions lost in the isoform (missing or distant residues); Blue: interactions conserved between the two forms; Green: new interactions detected specifically with iTRAIL-R4. Residues shown in red indicate those absent from the isoform due to alternative splicing. An asterisk (*) marks residues involved in interactions with both TRAIL subunits (interface). Interactions were identified from structural models visualized using PyMOL.

**Supplementary figure 3. Chemotherapeutic sensitization to TRAIL-induced cytotoxicity.** MDA-MB-231 cells, either wild-type (WT) or expressing canonical TRAIL-R4 (cTRAIL-R4) or iTRAIL-R4, were treated for 24 h with two concentrations of 5-fluorouracil (5FU), with increasing concentrations of TRAIL (0–1000 ng/mL) added during the final 12 h. Cell viability was assessed by MTT assay and expressed relative to untreated cells. Responses to TRAIL in combination with 5FU are presented in panels a-c, organized by 5FU concentration, and d-f, organized by TRAIL-R4 expression status (WT, canTRAIL-R4, and isoTRAIL-R4). Data represent mean ± SD of three independent technical replicates. Statistical analyses were performed using two-way ANOVA with multiple comparisons against iTRAIL-R4-expressing cells (*$p < 0.05$, **$p < 0.01$, ***$p < 0.001$, ****$p < 0.0001$).

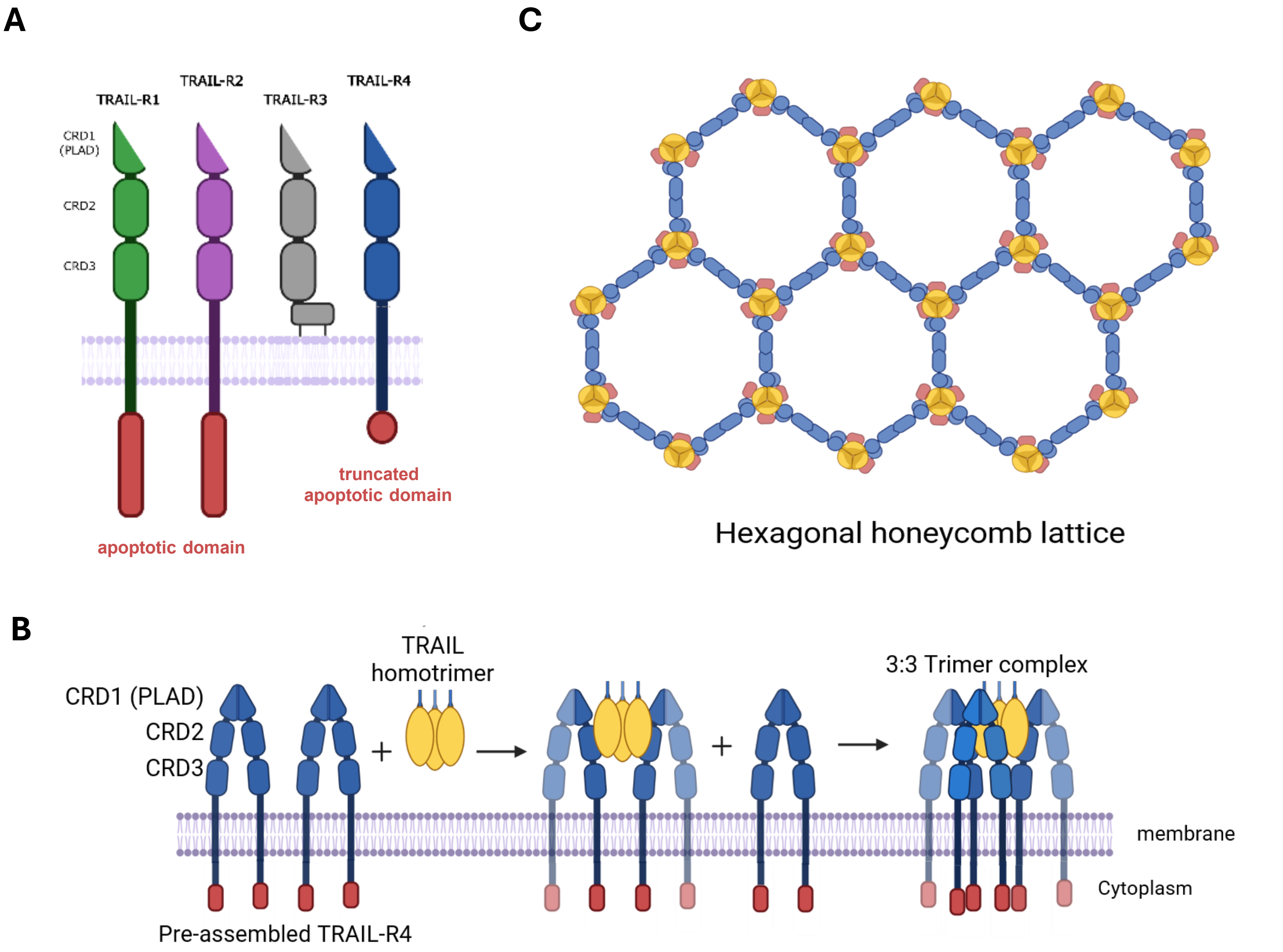


**Supplementary figure 1**

**A**

| Interaction Type | Zone | TRAIL Subunit | Residues TRAIL | Residues TRAIL-R4 | Distance (Å) |
|---|---|---|---|---|---|
| Ionic | B | α | R158 | E91 | 2,6 |
| | A | | D67 | K122 | 3,4 |
| | | β | K224 | E153 | 3,6 |
| | | | D203 | R156 | 4,8 |
| Polar | B | α | R158 | E91 | 3,2 |
| | A | | R130 | C118 | 3,2 |
| | | | | V120 | 2,9 |
| | | | D267 | K122 | 2,6 |
| | | | D269 | | 2,7 |
| | | | Y189 | E153* | 2,7 |
| | | | R191 | | 2,8-2,9 |
| | | | Q193 | S123 | 3,2 |
| | B | β | Y216 | I107 | 3 |
| | A | | K224 | E153* | 3,5-3,6 |
| | | | Q205 | | 2,8 |
| | | | | M154 | 2,8-3,1 |
| | | | N262 | N150 | 2,9-3,4 |
| | | | D203 | R156 | 2,7-2,8-3,3 |
| | | | K201 | | 3,1 |
| | | | | T157 | 2,7 |
| | | | N199 | R159 | 3,3 |

**D**

| Interaction Type | Zone | TRAIL Subunit | Residues TRAIL | Residues iTRAIL-R4 | Distance (Å) |
|---|---|---|---|---|---|
| Ionic | B | α | R158 | E91 | - |
| | A | | D267 | K122 | - |
| | | β | K224 | E153 | 2,2 |
| | | | D203 | R156 | 3,9 |
| Polar | B | α | R158 | E91 | - |
| | | | R158 | S77 | 3,6 |
| | | | G128 | K79 | 2,5 |
| | A | | R130 | C118 | - |
| | | | | V120 | - |
| | | | D267 | K122 | - |
| | | | D269 | | - |
| | | | Y189 | E153* | - |
| | | | R191 | E153* | 2,8 |
| | | | R191 | A85 | 3,5-3,5 |
| | | | Q193 | S123 | - |
| | | | Y237 | R156 | 3-3,4 |
| | | | E236 | T157 | 2,9 |
| | B | β | Y216 | I107 | - |
| | A | | K224 | E153* | 3,5-3,6 |
| | | | Q205 | | 2,8 |
| | | | | M154 | 2,8-3,1 |
| | | | N262 | N150 | 2,9-3,4 |
| | | | D203 | R156* | 2,4-3,0 |
| | | | K201 | | 3,1 |
| | | | K201 | T157 | - |
| | | | N199 | R159 | 5,6 |

**B**

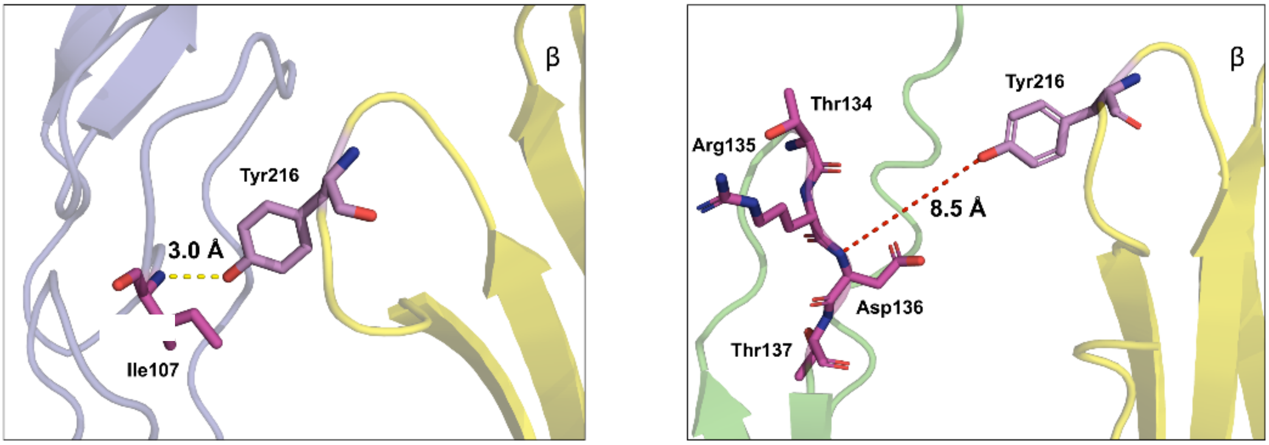


**C**

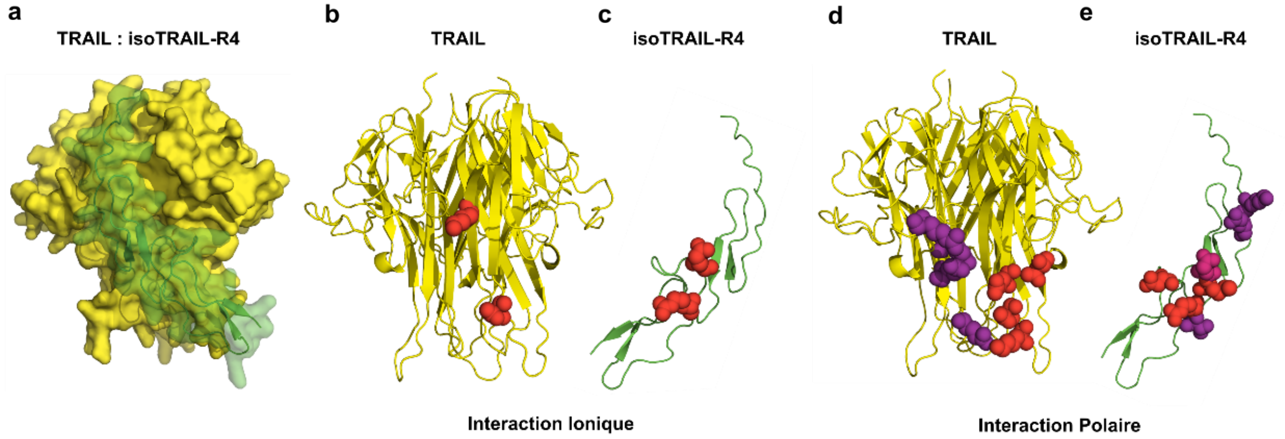


**Supplementary figure 2**

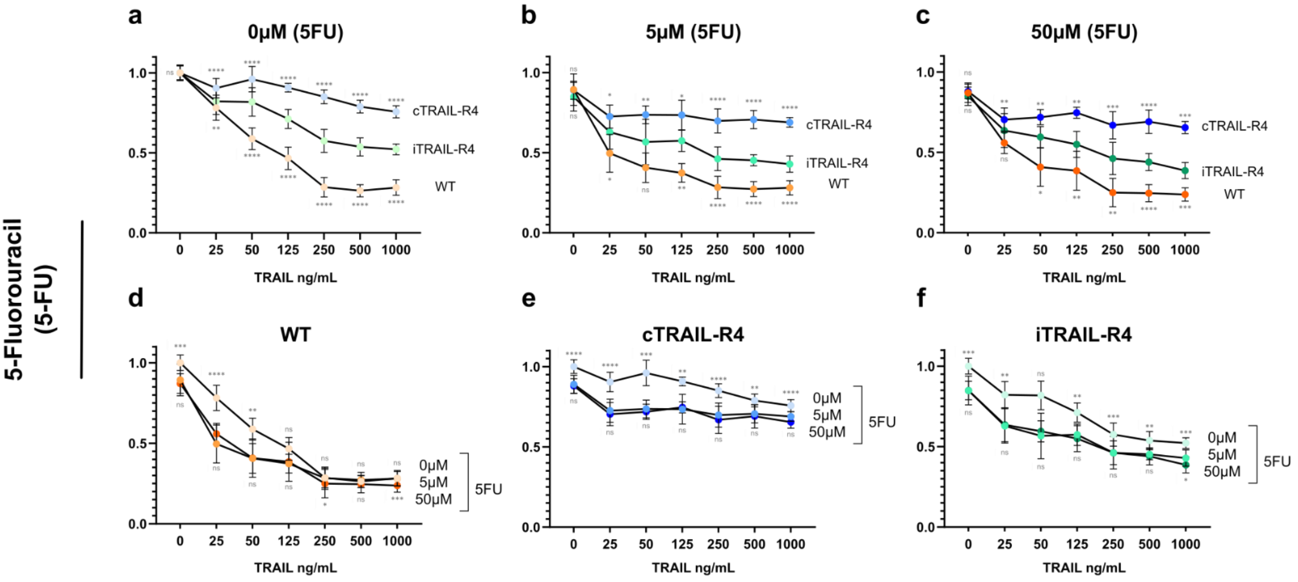


**Supplementary figure 3**
